# Implementation and Integration of Microbial Source Tracking in a River Watershed Monitoring Plan

**DOI:** 10.1101/514257

**Authors:** Elisenda Ballesté, Katalin Demeter, Bartholomew Masterson, Natàlia Timoneda, Wim G. Meijer

## Abstract

Fecal pollution of water bodies poses a serious threat for public health and ecosystems. Microbial source tracking (MST) using host specific bacteria are used to track the source of this potential pollution and be able to perform a better management of the pollution at the source. In this study we tested 12 molecular MST markers to track human, ruminant, sheep, horse, pig and gull pollution to determine their usefulness in their application for an effective management of water quality. First, the potential of the selected markers to track the source was evaluated using fresh fecal samples. Subsequently, we evaluated their performance in a catchment with different impacts, considering land use and environmental conditions. All MST markers showed high sensitivity and specificity, although none achieved 100% for both. Although some of the MST markers were detected in hosts other than the intended ones, their abundance in the target group was always several orders of magnitude higher than in the non-target hosts, demonstrating their suitability to distinguish between sources of pollution. The MST analysis matched the land use in the watershed allowing a very accurate assessment of the main hazards and sources of pollution, in this case mainly human and ruminant pollution. Correlating environmental parameters like temperature and rainfall with the levels of the MST markers provided insight into the dynamics of the pollution along the catchment. The levels of the human associated marker showed a significant negative correlation with rainfall in human polluted areas suggesting a dilution of the pollution, whereas at agricultural areas the ruminant marker increased with rainfall. There were no seasonal differences in the levels of human marker, indicating human pollution as a constant pressure throughout the year, whereas the levels of the ruminant marker was influenced by the seasons, being more abundant in summer and autumn. Performing MST analysis integrated with land uses and environmental data can improve the management of fecal polluted areas and set up good practices.

## Introduction

Fecal contamination of water bodies poses a serious threat as it introduces organic matter, nitrogen, phosphorous as well as potential pathogens causing health risks, environmental degradation, and economic losses, in particular when drinking water supplies and recreational and shellfish harvesting waters are affected [1]. A description of catchment characteristics, including possible sources of fecal pollution, is an essential prerequisite to facilitate efficient assessment, management and remediation of the affected region. The identification and assessment of causes of pollution that might affect bathing waters is enshrined in the European Bathing Water Directive 2006/7/EC [2].

A large number of library independent molecular markers to track the source of fecal pollution in water have been developed [3], which in general are based on the detection of host specific intestinal bacterial species using endpoint or quantitative PCR [4–11]. In comparison to other MST approaches, these library-independent techniques are cost effective, relatively easy to implement and provide fast results. DNA extracted from a single water sample can be used to analyze for the presence of multiple MST markers.

The composition of the animal and human intestinal microbiome is influenced by a number of factors, including age, diet and health status, as well as the geographical location of the host [12–14]. Host specific MST markers may therefore show a distinct geographical and temporal distribution [15,16]. It is therefore essential that MST markers are validated for the geographical area where they will be used. This includes determining the: i) abundance of MST markers at the source, since low initial concentration of these markers might become a problem when fecal matter is dispersed following its release in the environment, ii) the specificity of the marker, as it may be present in other hosts albeit at low concentrations and, iii) the temporal stability in different host groups [3]. It is therefore essential to establish these basic characteristics of the markers prior to their use in the field to assess water quality in a catchment.

The deployment of MST markers to assess water quality in a catchment is challenging, and a range of factors influencing the interpretation of the results of an MST analysis need to be considered. For example, fecal pollution in a catchment is frequently the result of contamination derived from a number of different sources, which increases the microbial diversity and complexity of a water sample. The variation in marker composition and abundance in a water body may introduce a bias in expected patterns. In addition, fecal pollution may arise from point or diffuse sources, which may be immediately and intermittently deposited in the waterbody as is often the case for wild life, settling on fields before it eventually enters a waterbody as runoff after rainfall, in for example agricultural pollution, or be continuously discharged from waste water treatment plants or through sewer misconnections, or sporadically from combined sewer overflows [17,18]. Spatial analysis using Geographical Information Systems (GIS) are becoming a very useful tool for water assessment [19]. Combining GIS with fecal indicator bacteria and MST, environmental data such as rainfall, flow and temperature may improve water management and assessment of specific areas [20,21].

In this study we have gone through several steps for the further implementation of MST markers as a toolbox for water routine monitoring. 1) We have evaluated the performance of different host specific markers detecting human, cow, sheep, horse, pig, and gull pollution on fecal samples collected within the study area. Sensitivity, specificity and abundance of the markers were addressed using PCR and real time quantitative PCR in these samples. 2) The MST markers showing a better performance were selected and tested in the study area, a river catchment with a potential source of animal and human fecal pollution addressed according to the land uses during a three year sampling period. 3) Finally the performance of the MST markers was validated using land cover and environmental (precipitation and temperature) data for the given catchment.

## Materials and Methods

### Study site

The Dargle River and its tributary rivers drain a catchment of approximately 133 km^2^ and flows into the Irish Sea in Bray, 27 km south of Dublin. The 20 km long river has a slope of approximately 2.7%, and a dry weather flow of 3 m^3^ s^−1^ that may increase 100-fold during severe weather events. There are extreme variations in climate within the catchment. In the eastern, coastal part of the catchment, annual rainfall is ca. 800 mm, there are approximately 185 days with rain per year, and monthly mean rainfall figures range between 50 mm in summer and 70 mm in winter. Mean daily temperatures lie between 5°C in winter and 15°C in summer; mean daily sunshine duration lies between 1.6 hours per day in winter and 5.6 hours in summer. In the mountains, rainfall is 1,500-2,000 mm per year, with snow-cover lasting a month; daily temperatures are much lower and sunshine hours fewer.

Although the Dargle catchment is relatively small (121 km^2^), it is characterized by a very diverse land use (Fig 1, Table 1). The lower catchment is mainly urban in nature, including the towns of Kilmacanogue, Enniskerry and Bray with 900, 2,700, and 32,000 inhabitants, respectively, while the upper catchment is characterized by tillage, cattle and sheep farming, forestry and peat bogs. The catchment is home to a number of riding stables.

**Table 1.** Percentage of the land uses surface affecting each sampling site in Dargle catchment. Locations and abbreviations of the sampling stations as in Fig 1.

|  | **DRG1** | **DRG2/3** | **KGH** | **KMG1** | **KMG2** | **CBR** | **SWN** | **GCU** | **GCR** |
| --- | --- | --- | --- | --- | --- | --- | --- | --- | --- |
| **Urban** | 0 | 0 | 0 | 0 | 1 | 2.4 | 96.2 | 4 | 0 |
| **Sport** | 0 | 10.1 | 0 | 3.1 | 2.1 | 26.1 | 0 | 2.3 | 0 |
| **Agriculture** | 3 | 52.5 | 70.6 | 55.1 | 59.1 | 60.1 | 2.2 | 36.8 | 14.4 |
| **Forest** | 31.6 | 33.1 | 5.3 | 1.7 | 6.8 | 11.4 | 1.3 | 26.4 | 35.5 |
| **Peatland** | 65.4 | 4.3 | 24.1 | 40.1 | 31 | 0 | 0.3 | 30.5 | 48.8 |
| **Lakes** | 0 | 0 | 0 | 0 | 0 | 0 | 0 | 0 | 1.3 |

**Figure 1.**
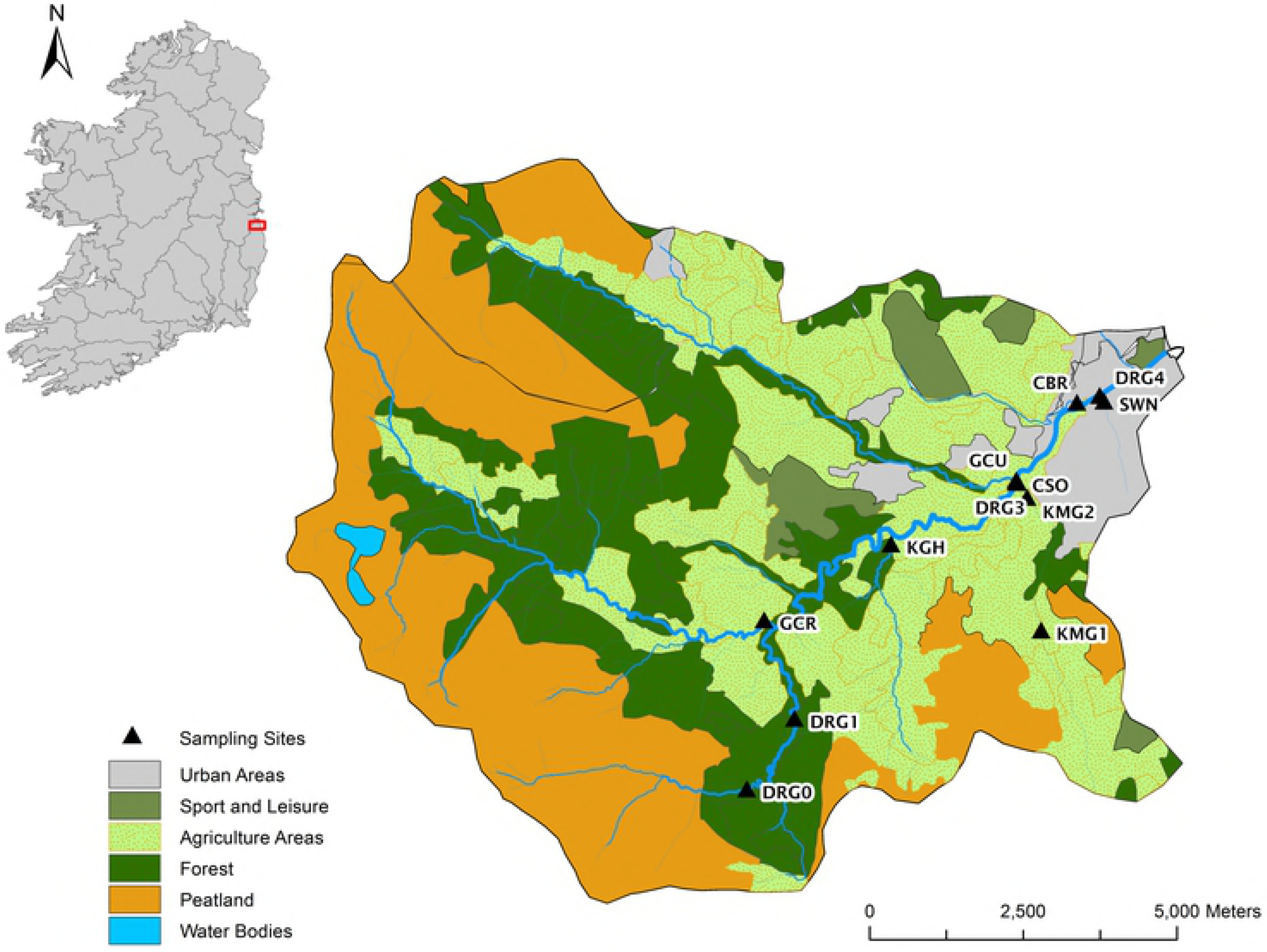
Dargle catchment land uses and sampling sites. Location of the sampling sites and land uses of the Dargle river catchment.

### Sample Collection

Water samples (n=354) were collected over a period of 3 years at 10 sampling sites, located along the Dargle river (DRG1, DRG2 and DRG3) and in the main tributaries: Swan (SWN), County Brook (CBR), Kilmacanogue (KMG1 and KMG2), Killough (KGH), Glencullen (GCU) and Glencree (GCR) rivers (Figure 1). These sampling sites were selected to be representative of the different land uses and tributaries. Sewage effluent samples (n=25) were collected from the Enniskerry waste water treatment plant (6,000 p.e.) (secondary treatment with nutrient reduction) before discharging into the Dargle River. Samples were collected in sterile containers, transported on ice and stored at 4°C until analysis.

Fresh fecal samples (n=181) were collected from pigs (n=20), horses (n=27), sheep (n=25), cattle (n=22), deer (n=15), goats (n=2), peacock (n=3) and a donkey (n=1). Samples from gulls (n=32) and swans (n=8) were collected in Bray harbor and Irelands’ Eye island. The main gull species in these environments are great black-backed gull (*Larus marinus*), herring (*Larus argentatus*) and common gulls (*Larus canus*). Samples of each animal source were collected from at least two different locations. Human samples (n=26) were obtained from Crumlin Hospital (Dublin, Ireland) from individuals not receiving antibiotic treatment.

### Enumeration of FIB

The levels of total coliforms, *E. coli* and intestinal enterococci were determined using Colilert-18 and Enterolert with a Quanti-Tray/2000 system (IDEXX Laboratories, UK) according to manufacturer’s instructions.

### DNA extraction

DNA was extracted from water samples following filtering of 100 ml water using a filter with 0.2 μm pore-size (Supor 200 PES, Pall Corporation, NY). Filters were placed in 0.5 ml of GITC buffer (5 M guanidine thiocyanate, 100 mM EDTA [pH 8.0], 0.5% Sarkosyl) and frozen at −20°C in GITC buffer until DNA extraction. DNA was extracted using the DNeasy Blood & Tissue Kit (Qiagen) with some modifications as reported previously [22].

DNA was extracted from 180-250 mg of fecal sample using the QIAamp DNA Stool MiniKit (Qiagen, UK) according to the manufacturer’s instructions.

### Detection of MST markers by end-point PCR and real time quantitative PCR

#### End-point PCR

Detection of the MST markers by PCR was performed using primers described previously (S1 Table). Five host-specific markers matched the 16S rRNA gene of *Bacteroidales* species: the human marker HF183, the ruminant marker CF183 [4], the horse marker HoF597 and the pig markers PF163 [23] and Pig-2-Bac [24].

The MST markers were amplified in a 25-μl PCR mixture containing Green GoTaq Flexi Buffer (Promega), 0.2 μM of each primer, 0.625 U GoTaq DNA polymerase (Promega), 2 mM MgCl_2_, 200 μM dNTP and 0.1 mg/ml nonacetylated BSA (Applied Biosystems). The reaction mixture was incubated at 95°C for 2 min, followed by 35 (fecal sample) or 45 (water sample) cycles of 94°C for 30s, an appropriate annealing temperature for each marker (Table S1) for 30 s, and an extension step at 72°C for 20 s (HoF597 and Pig-2-Bac marker) or 40 s (HF183, CF128 and PF163); followed by a final 5 min extension at 72°C.

The 16S rRNA gene of *Catellicoccus marimammalium* was amplified using primers GullF and GullR (S1 Table) [9]. The reaction mixture was incubated at 95°C for 2 min followed by 35 cycles of 94°C for 30 s, 63°C for 30 s, and 72°C for 35 s, which were followed by a final incubation at 72°C for 5 min.

Nested-PCR was used to detect porcine and ovine mitochondrial DNA [25] (Table S1). The reaction mixtures were incubated at 95°C for 2 min, followed by 35 cycles consisting of 94°C for 40 s, 55°C for 50 s, and at 72°C for 45 s; followed by a final 5 min extension.

The universal primers F63 and 1389R [26,27] were used (0.4 μM) to amplify the 16S rRNA gene, and was used as a positive control to rule out the presence of PCR inhibitors in samples causing false negative results. The reaction mixture was incubated at 95°C for 2 min, followed by 30 cycles of 94°C for 45 s, 55°C for 45 s, and 72°C for 90 s, followed by 72°C for 5 min. Controls included filtration blanks, DNA extraction blanks, no-template PCR control and PCR positive control.

#### Real Time Quantitative PCR

The forward primers HF183F and CF128F were paired with the general *Bacteroidales* primer 265R and HoF597F with Bac708R [4,23,28] (S1 Table). In addition, a CowM3 marker matching a specific gene (sialic acid-specific 9-*O*-acetylesterase secretory protein homologue) from cow-specific *Bacteroidales* was tested [29]. The levels of general *Bacteroidales* were quantified using AllBac-296R and AllBac-412R primers [8] (Table S1). The assays were performed using a Light Cycler 480 Instrument (Roche Diagnostics, Mannheim, Germany). Reaction mixtures (10 μl) consisted of DNA Master SYBR Green I Buffer (Roche) and 0.5 μM of each primer. The amplification conditions were: an initial denaturation for 10 min at 95°C, followed by 45 cycles at 95°C for 15 s, 60°C for 15 s and an extension step at 72°C for 15 s. A melting curve analysis was carried out by heating the PCR products to 95°C, cooled to 60°C for 15 s and slowly reheated to 95°C at a rate of 0.1 °C/s. PCR conditions to amplify and quantify CowM3 marker were as described [29]. Samples were run in triplicate with 1:10 fold dilution of the samples in order to detect potential inhibition effects. The results were expressed as gene copies (gc) 100 mg^−1^ or 100 ml^−1^. Controls included no-template PCR control and PCR positive control. The detection limit was established through the analysis of serial dilution of fecal and water samples. The quantification limits were 200 gc 100 ml^−1^ for qHF183, qHoF597, CowM3 and AllBac, and in the case of the CF128 marker, the quantification limit was higher at 1,000 gc 100 ml ^−1^. Lower concentrations could be detected but not quantified accurately due to the variability when working with low concentration of DNA.

### Quantification of Land Use

ArcGIS 9.3 was used to visualize and analyze CORINE land cover data (CLC2006, http://gis.epa.ie) in order to assess different land use in the Dargle catchment. Additional GIS data was obtained from the Ordnance Survey Ireland (OSi). These layers included river segments, catchment and subcatchments areas and digital elevation models.

### Statistical Analysis

Sensitivity and specificity for the eight microbial source tracking markers were calculated as described previously [30]. In order to obtain a 95% confidence interval for the estimates of sensitivity and specificity, at least 20 fecal samples of target species were analyzed. Sensitivity (r) and specificity (s) are defined as r=a/(a+c) and s=d/(b+d), where a is when a fecal DNA sample is positive for the PCR marker of its own target (true positive); b is when a fecal DNA sample is positive for a PCR marker of another target (false positive); c is when a fecal DNA sample is negative for a PCR marker of its own target (false negative); and d is when a fecal DNA sample is negative for a PCR marker of another target (true negative).

The positive predictive value or conditional probability that a particular source of fecal contamination was present when a water sample tested positive for the corresponding MST marker was calculated using Bayes’ Theorem [7].

The quantitative values of the qPCR markers were log_10_ converted to achieve normality. Despite of this, not all the variables were normally distributed, thus, Spearman correlation was used to detect relationships among variables. Data was analyzed using R, in the RStudio interface, version 0.97.314, R version 3.1.2 (2014-10-31)—“Pumpkin Helmet” [31]. Other primary packages, in addition to base packages, used in this analysis included *car*, *agricolae* and *ggplot2* [32–34].

### Ethics Statement

This research was not subject to ethical approval. No permits were required for the described study, which complied with all relevant regulations.

## RESULTS

### Sensitivity and specificity of host-specific markers in feces and sewage

It was shown previously that significant geographical variability exists in the sensitivity and specificity of MST markers [15,30]. The performance of these MST markers was therefore evaluated using feces from local animals and humans, prior to deploying these in the field. DNA was extracted from fresh fecal samples from mammals: cattle, deer, donkey, goat, human, horse, pig, sheep, and from birds: gull, peacock, and swan. Initially the general 16S rRNA gene was amplified, showing that most samples contained amplifiable DNA, except for DNA extracted from 6 out of 32 gull samples that could not be amplified and were discarded.

The performance of the MST markers was very good, achieving a sensitivity higher than 85% for most of the markers. In the case of the human marker HF183, a sensitivity of 73% was obtained when individual fecal samples were analyzed, however, the sensitivity of the human marker was 100% using raw and treated sewage samples. The specificities of all markers were 87% or higher (Table 1).

**Table 1.**
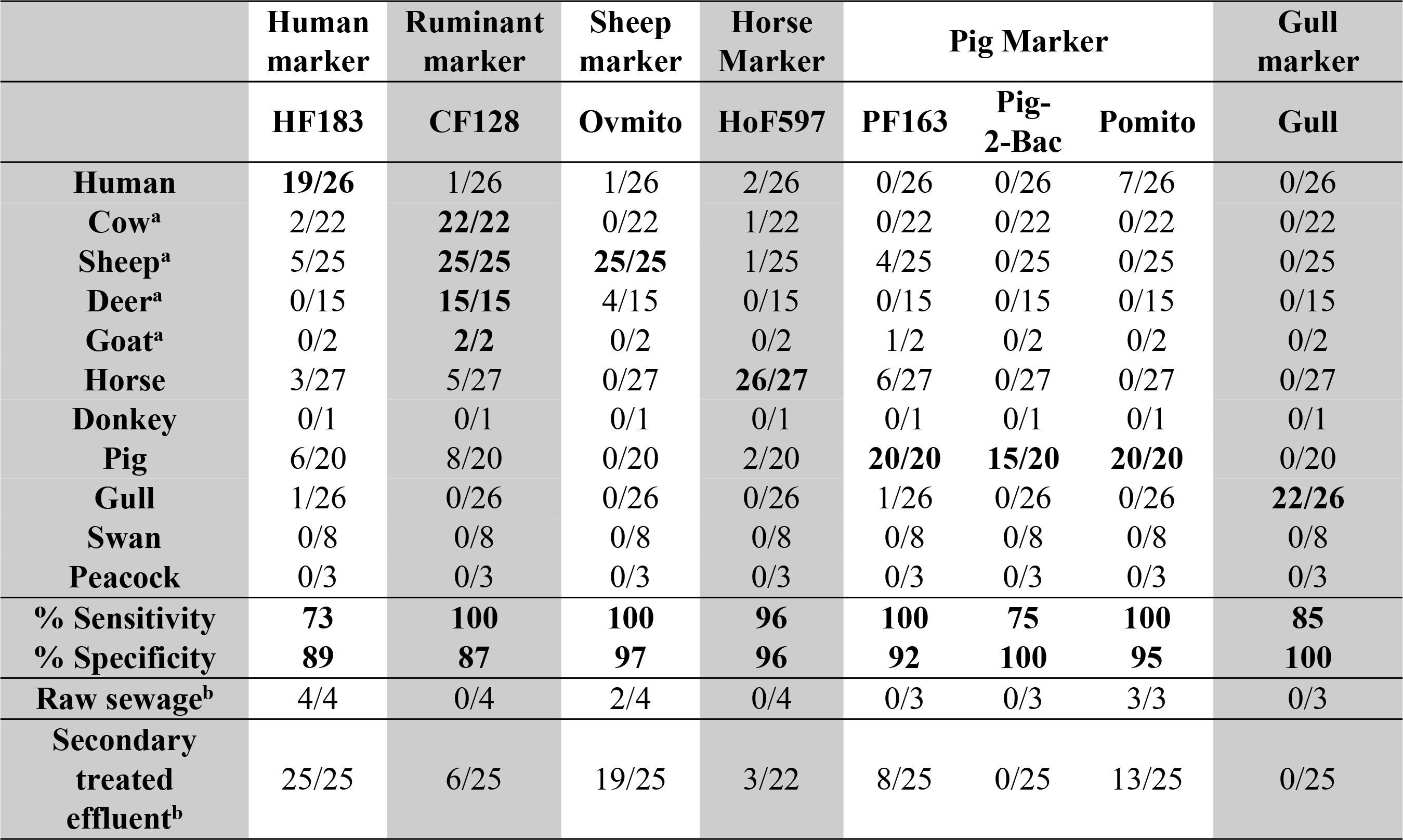
Number of positive samples, sensitivity and specificity for different MST markers evaluated by PCR in fecal samples.

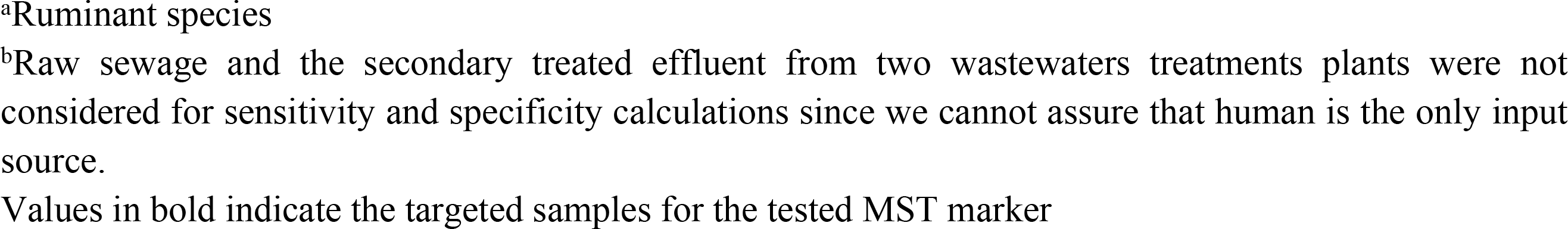

### MST levels of host-specific markers in feces and sewage by qPCR

The levels of human (HF183), ruminant (CF128), cow (CowM3) and horse (HoF597) markers and total *Bacteroidales* (AllBac) were measured by qPCR in fecal samples from different sources (Figure 2, Table 2). The number of positive samples, as determined by qPCR, was higher than by end-point PCR. The samples that were positive for qPCR but negative for endpoint PCR contained very low target DNA concentrations and in many cases were below the quantification limit.

**Table 2.**
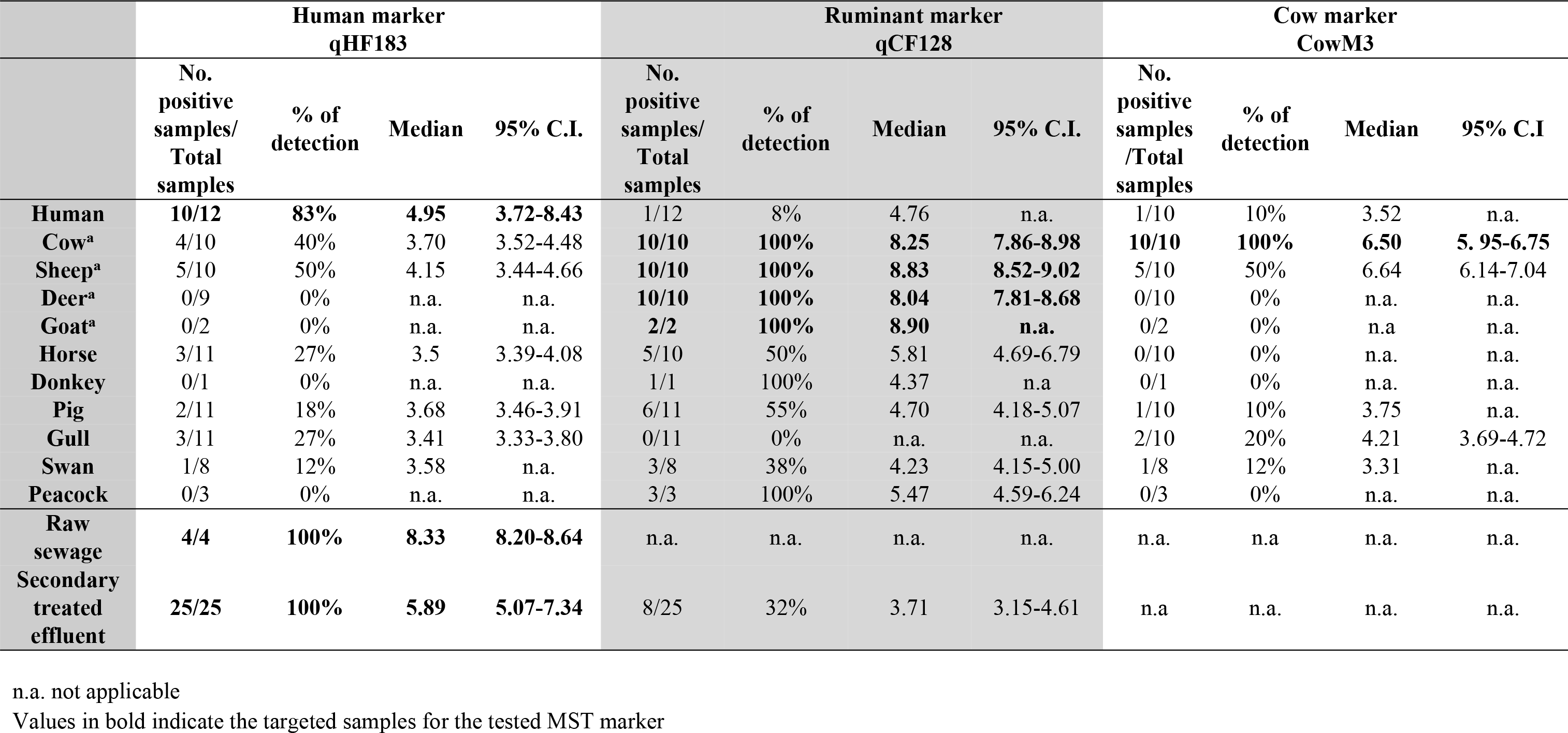
Positive samples, percentage of detection, median and 95% confidence interval (C.I.) of the log_10_ transformed levels of the MST markers (gc 100 mg^−1^ or gc 100 ml-^1^) evaluated by qPCR in fecal and wastewater samples.

|  | Human marker<br>qHF183 |  |  |  | Ruminant marker<br>qCF128 |  |  |  | Cow marker<br>CowM3 |  |  |  |
| --- | --- | --- | --- | --- | --- | --- | --- | --- | --- | --- | --- | --- |
|  | No.<br>positive<br>samples/<br>Total<br>samples | % of<br>detection | Median | 95% C.I. | No.<br>positive<br>samples/<br>Total<br>samples | % of<br>detection | Median | 95% C.I. | No.<br>positive<br>samples<br>/Total<br>samples | % of<br>detection | Median | 95% C.I. |
| **Human** | **10/12** | **83%** | **4.95** | **3.72-8.43** | 1/12 | 8% | 4.76 | n.a. | 1/10 | 10% | 3.52 | n.a. |
| **Cow<sup>a</sup>** | 4/10 | 40% | 3.70 | 3.52-4.48 | **10/10** | **100%** | **8.25** | **7.86-8.98** | **10/10** | **100%** | **6.50** | **5.95-6.75** |
| **Sheep<sup>a</sup>** | 5/10 | 50% | 4.15 | 3.44-4.66 | **10/10** | **100%** | **8.83** | **8.52-9.02** | 5/10 | 50% | 6.64 | 6.14-7.04 |
| **Deer<sup>a</sup>** | 0/9 | 0% | n.a. | n.a. | **10/10** | **100%** | **8.04** | **7.81-8.68** | 0/10 | 0% | n.a. | n.a. |
| **Goat<sup>a</sup>** | 0/2 | 0% | n.a. | n.a. | **2/2** | **100%** | **8.90** | **n.a.** | 0/2 | 0% | n.a. | n.a. |
| **Horse** | 3/11 | 27% | 3.5 | 3.39-4.08 | 5/10 | 50% | 5.81 | 4.69-6.79 | 0/10 | 0% | n.a. | n.a. |
| **Donkey** | 0/1 | 0% | n.a. | n.a. | 1/1 | 100% | 4.37 | n.a. | 0/1 | 0% | n.a. | n.a. |
| **Pig** | 2/11 | 18% | 3.68 | 3.46-3.91 | 6/11 | 55% | 4.70 | 4.18-5.07 | 1/10 | 10% | 3.75 | n.a. |
| **Gull** | 3/11 | 27% | 3.41 | 3.33-3.80 | 0/11 | 0% | n.a. | n.a. | 2/10 | 20% | 4.21 | 3.69-4.72 |
| **Swan** | 1/8 | 12% | 3.58 | n.a. | 3/8 | 38% | 4.23 | 4.15-5.00 | 1/8 | 12% | 3.31 | n.a. |
| **Peacock** | 0/3 | 0% | n.a. | n.a. | 3/3 | 100% | 5.47 | 4.59-6.24 | 0/3 | 0% | n.a. | n.a. |
| **Raw**<br>**sewage** | **4/4** | **100%** | **8.33** | **8.20-8.64** | n.a. | n.a. | n.a. | n.a. | n.a. | n.a. | n.a. | n.a. |
| **Secondary**<br>**treated**<br>**effluent** | **25/25** | **100%** | **5.89** | **5.07-7.34** | 8/25 | 32% | 3.71 | 3.15-4.61 | n.a. | n.a. | n.a. | n.a. |

n.a. not applicable
Values in bold indicate the targeted samples for the tested MST marker

n.a. not applicable Values in bold indicate the targeted samples for the tested MST marker
|  | **Horse Marker**<br>**qHoF597** |  |  |  | ***Bacteroidales* marker**<br>**AllBac** |  |  |  |
| --- | --- | --- | --- | --- | --- | --- | --- | --- |
|  | **No.**<br>**positive**<br>**samples/**<br>**Total**<br>**samples** | **% of**<br>**detection** | **Median** | **95% C.I** | **No.**<br>**positive**<br>**samples/**<br>**Total**<br>**samples** | **% of**<br>**detection** | **Median** | **95% C.I** |
| **Human** | 1/11 | 9% | 3.68 | n.a. | **12/12** | **100%** | **8.76** | **5.95-9.71** |
| **Cow<sup>a</sup>** | 5/10 | 50% | 4.36 | 3.39-4.89 | **10/10** | **100%** | **8.62** | **8.48-9.02** |
| **Sheep<sup>a</sup>** | 6/10 | 60% | 5.00 | 3.64-5.24 | **10/10** | **100%** | **9.09** | **8.66-9.29** |
| **Deer<sup>a</sup>** | 4/10 | 40% | 3.74 | 3.41-3.97 | **10/10** | **100%** | **8.03** | **7.85-8.58** |
| **Goat<sup>a</sup>** | 2/2 | 100% | 4.28 | n.a. | **2/2** | **100%** | **8.72** | **n.a.** |
| **Horse** | **10/10** | **100%** | **6.32** | **4.94-6.99** | **11/11** | **100%** | **8.92** | **8.36-9.19** |
| **Donkey** | 1/1 | 100% | 4.83 | n.a. | **1/1** | **100%** | **7.95** | **n.a.** |
| **Pig** | 6/11 | 55% | 4.98 | 4.53-5.40 | **10/10** | **100%** | **9.42** | **9.14-9.78** |
| **Gull** | 2/11 | 18% | 4.04 | n.a. | **11/11** | **100%** | **4.00** | **3.36-5.83** |
| **Swan** | 0/8 | 0% | n.a. | n.a. | **8/8** | **100%** | **5.65** | **4.06-6.80** |
| **Peacock** | 0/3 | 0% | n.a. | n.a. | **3/3** | **100%** | **8.55** | **7.75-9.01** |
| **Raw**<br>**sewage** | n.a. | n.a. | n.a. | n.a. | **4/4** | **100%** | **9.92** | **9.80-10** |
| **Secondary**<br>**treatment**<br>**effluent** | 4/20 | 20% | 3.67 | 3.22-4.63 | **23/23** | **100%** | **7.35** | **6.97-8.17** |

**Figure 2.**
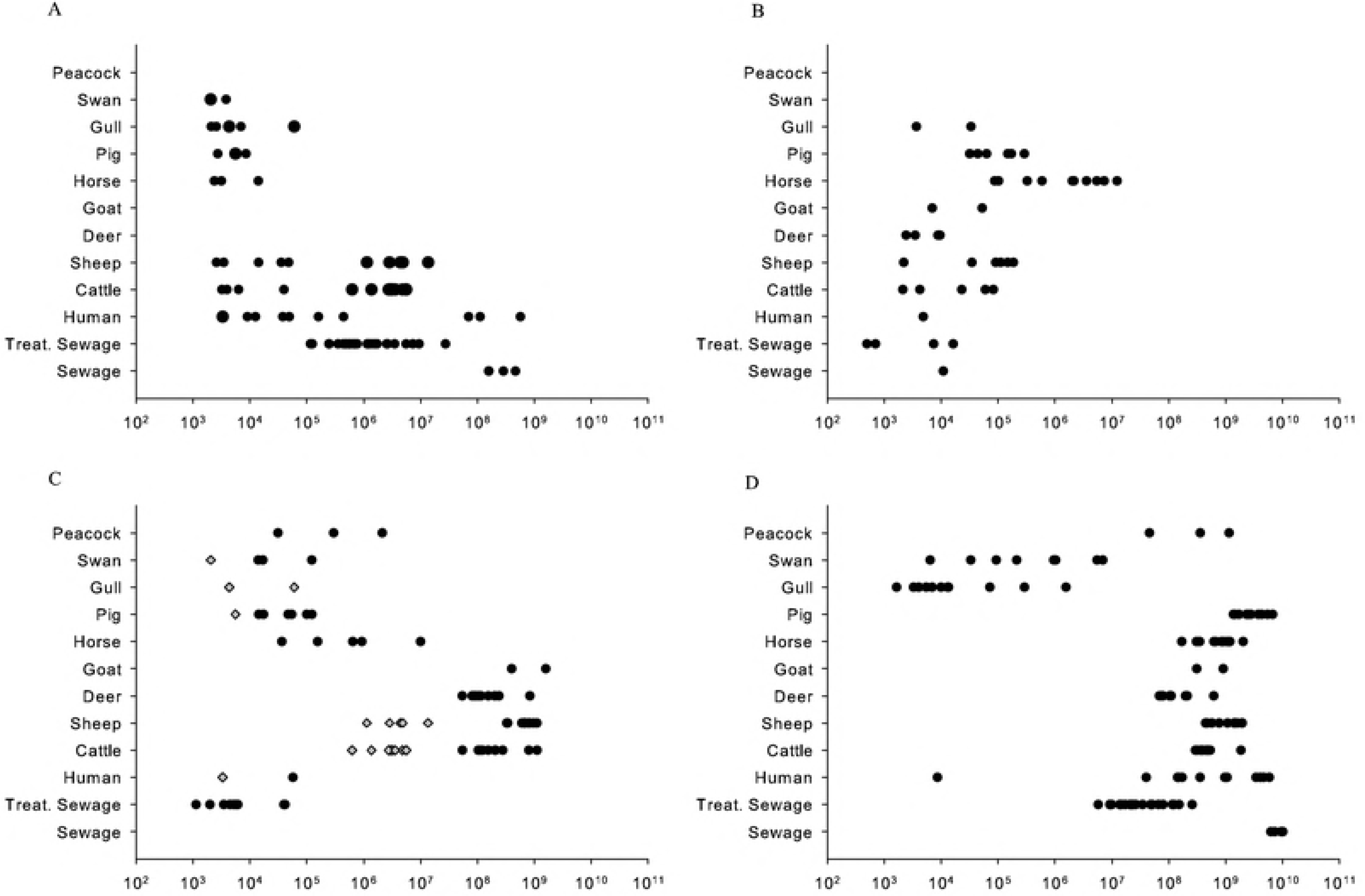
Abundance of the different MST markers in fecal samples from different animal sources. (A) Human marker (HF183), (B) Horse Marker (HoF597), (C) Ruminant Marker (CF138) (in black) and Cow Marker (CowM3) (in grey) and (D) General *Bacteroidales* (AllBac). Values are expressed in gene copies 100 mg^−1^ or gc ml^−1^.

The HF183 marker displayed considerable variability in individual human samples, ranging from 3.7 to 8.4 log_10_ gc/100 mg (Figure 2A). As expected, there was less variability and higher average levels of human marker in raw and treated sewage. The concentration of the horse (HoF597) marker was an order of magnitude (1.3 to 2.5 log_10_) higher in horse feces than in feces from other animal sources (Figure 2B). The ruminant (CF128) marker levels were around 2.5 − 4.0 log_10_ higher in ruminant samples compared to feces from humans and other animal sources (Figure 2C). In contrast to the human marker, the ruminant marker displayed low variability among individuals of the same group. The levels of the cow marker (CowM3) were 2.3 − 3.2 log_10_ higher in feces from cows and sheep than from other sources. Surprisingly, half of the sheep samples were positive for this marker. The levels of general *Bacteroidales* were high and similar for all the tested mammalian fecal samples. In contrast, the fecal samples from swans and gulls showed lower levels of the general *Bacteroidales* marker. The negative controls for sample processing, nucleic DNA extraction, and PCR and qPCR reactions were in all cases negative.

To verify whether significant differences existed when comparing qualitative end-point PCR and the levels of host-marker levels among the corresponding target and non-target samples, as determined by qPCR, a Chi-squared and independent sample Kruskal-Wallis Test analysis was used. The differences within the target and non-targeted samples were statistically significant for all the markers evaluated in the study (P < 0.05). Thus, the MST markers tested are potentially useful to discern among different fecal samples in Ireland.

### Extreme variation in fecal contamination is related to land use

The Dargle catchment is characterized by a highly varied land use, including pristine, agricultural, forested and urban areas (Fig 1), which were analyzed using Arc Hydro tools of ArcGIS using CORINE land cover data. This informed the localization of sampling stations within the catchment, with a view to capture the impact of land use on water quality. The catchment was sampled at 10 stations (Fig 1). In addition to MST markers, the levels of fecal indicator bacteria (FIB), i.e. *E. coli* and Enterococci, were determined to evaluate the level of fecal pollution in the sub-catchments of the Dargle.

The highest levels of FIB (Fig 3A, Fig 3B) in the Dargle catchment over a 3 year period were obtained from sampling stations surrounded by urban areas: The Swan (SWN) and Kilmacanogue (KMG2) rivers, and the Dargle River after the confluence of the outfall (CSO) of the Enniskerry WWTP (DRG3). The DRG3 station was located just 30 m downstream DRG2, however the secondary treatment effluent of the WWTP discharged between the two points explaining an increase of one order of magnitude in the *E. coli* and Enterococci levels (Fig 3A, 3B). FIB levels were lower upstream in the catchment. Interestingly, there was a 10-fold increase in FIB levels in the Kilmacanogue River when comparing the sampling sites KMG1 and KMG2, which are separated by approximately 2 km. Lower levels were detected in County Brook (CBR), Killough River (KGH), Kilmacanogue River upstream of Kilmacanogue town (KMG1), and the Dargle before the Enniskerry WWTP outfall (DRG2). Pristine waters are found in the Glencullen (GCU) and Glencree (GCR) Rivers, and the upstream reaches of the Dargle River (DRG1).

**Figure 3.**
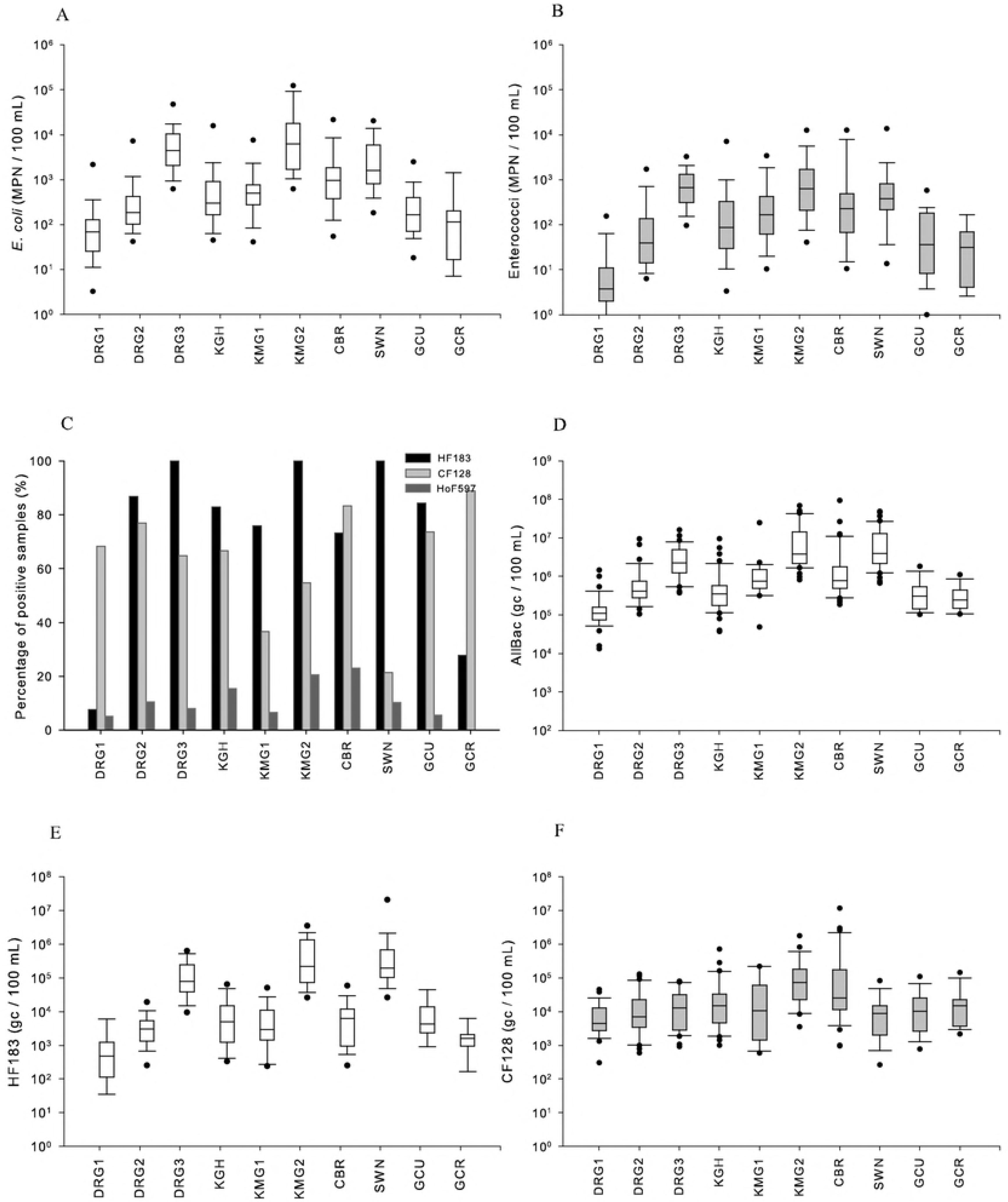
Levels of the fecal indicators and MST markers along the Dargle catchment. (A) *E. coli* (MPN 100 ml^−1^), (B) Enterococci (MPN 100 ml^−1^), (C) Percentage of positive samples detected by PCR for the HF183, CF128 and HoF597 markers, (D) levels of AllBac, (E) HF183 and (F) CF128 by qPCR (gc 100 ml^−1^). Box of the box plots represents the 25^th^, 50^th^ and 75^th^ percentile, the whiskers the 10% and 90^th^ percentile and the dots the 5th and 95^th^ percentile. Locations and abbreviations of the sampling stations as in Fig 1.

### Land use and the biological source of fecal water pollution

The Dargle catchment is very diverse in the levels of fecal contamination (Fig 3, Table 3), which is to be expected based on its land use (Figure 1). In order to analyze the biological sources of pollution, water samples were initially analyzed using end-point PCR. The presence/absence of the human (HF183), ruminant (CF128) and horse (HoF597) MST markers were evaluated by end-point PCR in the Dargle catchment, since these target the main pollution pressures in the areas of interest (human, ruminant and equine).

**Table 3.** Presence and abundance of the FIB and MST markers in freshwater samples. Median and 95% confidence interval (C.I.) of the log_10_ transformed data from the MST markers evaluated by qPCR. Positive samples / Total samples (PS/TS), percentage of positive samples evaluated by PCR. Locations and abbreviations of the sampling stations as in Fig 1.

|  | DRG1 |  | DRG2 |  | DRG3 |  | KGH |  | KMG1 |  |
| --- | --- | --- | --- | --- | --- | --- | --- | --- | --- | --- |
|  | Median | 95% C.I. | Median | 95% C.I. | Median | 95% C.I. | Median | 95% C.I. | Median | 95% C.I. |
| *E. coli* | 1.84 | 0.64-2.71 | 2.27 | 1.72-3.36 | 3.65 | 2.84-4.36 | 2.48 | 1.71-3.85 | 2.70 | 1.84-3.47 |
| Ent | 0.57 | <1-1.94 | 1.59 | 0.80-3.16 | 2.83 | 2.05-3.40 | 1.94 | 0.73-3.55 | 2.22 | 1.18-3.29 |
| qHF183 | 2.68 | 1.61-3.67 | 3.48 | 2.48-4.15 | 4.89 | 4.07-5.79 | 3.70 | 2.56-4.71 | 3.47 | 2.41-4.57 |
| qCF128 | 3.65 | 3.16-4.52 | 3.84 | 2.96-5.01 | 4.11 | 3.16-4.89 | 4.18 | 3.20-5.36 | 4.02 | 2.84-5.33 |
| AllBac | 5.05 | 4.53-5.80 | 5.62 | 5.13-6.55 | 6.35 | 5.70-6.97 | 5.55 | 4.90-6.58 | 5.88 | 5.50-6.34 |
|  | PS/TS | % of positive | PS/TS | % of positive | PS/TS | % of positive | PS/TS | % of positive | PS/TS | % of positive |
| HF | 3/40 | 7.5% | 33/38 | 86.8% | 36/36 | 100% | 34/41 | 82.9% | 22/29 | 75.9% |
| CF | 28/41 | 68.3% | 30/39 | 76.9% | 24/37 | 64.9% | 28/42 | 66.7% | 11/30 | 36.7% |
| HoF | 2/39 | 5.1% | 4/38 | 10.5% | 3/37 | 8.1% | 6/39 | 15.4% | 2/30 | 6.7% |

|  | KMG2 |  | CBR |  | SWN |  | GCU |  | GCR |  |
| --- | --- | --- | --- | --- | --- | --- | --- | --- | --- | --- |
|  | Median | 95% C.I. | Median | 95% C.I. | Median | 95% C.I. | Median | 95% C.I. | Median | 95% C.I. |
| *E. coli* | 3.80 | 2.81-5.03 | 2.98 | 1.97-3.99 | 3.20 | 2.26-4.31 | 2.22 | 1.67-3.02 | 2.06 | 0.85-3.17 |
| Ent | 2.81 | 1.72-4.00 | 2.34 | 1.04-4.04 | 2.59 | 1.13-4.14 | 1.56 | 0.54-2.44 | 1.49 | 0.41-2.23 |
| qHF183 | 5.35 | 4.44-6.35 | 3.81 | 2.64-4.62 | 5.29 | 4.42-7.32 | 3.63 | 2.96-4.66 | 3.20 | 2.38-3.67 |
| qCF128 | 4.87 | 3.93-5.88 | 4.41 | 3.37-6.44 | 3.94 | 2.42-4.92 | 4.01 | 3.13-4.79 | 4.18 | 3.48-4.98 |
| AllBac | 6.59 | 6.07-7.66 | 5.89 | 5.38-7.13 | 6.60 | 5.88-7.66 | 5.49 | 5.06-6.14 | 5.38 | 5.03-5.93 |
|  | PS/TS | % of positive | PS/TS | % of positive | PS/TS | % of positive | PS/TS | % of positive | PS/TS | % of positive |
| HF | 41/41 | 100% | 30/41 | 73.2% | 41/41 | 100% | 16/19 | 84.2% | 5/18 | 27.8% |
| CF | 23/42 | 54.8% | 35/42 | 83.3% | 9/42 | 21.4% | 14/19 | 73.7% | 16/18 | 88.9% |
| HoF | 8/39 | 20.5% | 9/39 | 23.1% | 3/39 | 10.3% | 1/18 | 0% | 0/18 | 0% |

The human marker was detected in all samples collected at the DRG3, KMG2 and SWN sampling stations (Figure 3C). The DRG3 site is impacted by the effluent from the Enniskerry WWTP, which is located 30 meters upstream of the sampling site. The KMG2 site is located in Kilmacanogue, downstream of Kilmacanogue town. The Swan River traverses several housing estates in Bray. The human marker was also detected with high frequency (more than 80% of positive samples) in DRG2, KGH and GCU, but was virtually absent in samples taken at the DRG1 station (<10%), which is located in the pristine areas of the Dargle catchment (Figure 3C, Table 3). The ruminant marker was less prevalent than the human marker in water samples. CF183 was detected in more than 60% of the sampling stations surrounded by agricultural areas: DRG1, DRG2, DRG3, KGH, CBR, GCU and GCR, but was virtually absent in the SWN (~20%), which flows through an almost exclusive urban area (Figure 3F, Table 3). While the horse marker (HoF597) was rarely observed in general, its presence stood out at the KMG2 and CBR sampling stations (more than 20% of samples tested positive). There are several horse riding stables located upstream from these sampling sites.

Bayesian statistics was used to determine the probability of detecting feces from the target-host within a given station using the end-point PCR results. Given the prevalence of the three MST markers in the catchment, as well as the sensitivity and specificity of these markers as determined using fecal samples (Table 1), the conditional probability of detecting the biological source of pollution was evaluated [7]. The conditional probability of having human pollution when the HF183 marker is detected was 0.95. The conditional probabilities for the ruminant marker (CF128) and horse marker (HoF597) were 0.93 and 0.75 respectively.

The levels of the human MST marker (qHF183) were high (10^5^-10^6^ gc 100 ml^−1^) in sampling sites with anthropogenic impact: SWN and KMG2, DRG3 and the CSO. Intermediate levels of the human marker, between 10^3^-10^4^ gc 100 ml^−1^, were detected in sampling sites with medium anthropogenic impact: KGH, CBR, GCU, KMG1 and DRG2. The levels of the human marker were below or at the detection limit at DRG1 and GCR (Fig 3E and Table 3).

There was a fairly even distribution of the levels of the ruminant marker along the different sampling points of the Dargle catchment (Fig 3F and Table 3). However, in sampling points with an agricultural impact: KGH, KMG1, KMG2 and CBR, the levels of the ruminant marker soared sporadically, often related to rainfall or unknown events.

### Correlation between MST marker and FIB levels

The level of general *Bacteroidales* marker (using AllBac marker) was assessed in order to evaluate the potential use of total *Bacteroidales* as a fecal indicator. The levels of AllBac were higher than 10^4^ gc 100 ml^−1^ in all samples (Fig 3D) and they correlated with the levels of *E. coli* (r_s_: 0.797; n: 354) and Enterococci (r_s_: 0.687; n: 354). With respect to individual sampling stations, AllBac correlated with the *E. coli* levels at all stations, expect for the upstream sampling sites: DRG1 and GCR, which displayed none or very low levels of *E. coli* and Enterococci. Only five of the sampling sites showed a significant correlation between the AllBac marker and Enterococcus levels (Table S2).

The human marker (qHF183) levels were strongly correlated with those of *E. coli* and Enterococci and AllBac in KMG2 (r_s_ = 0.666; r_s_ = 0.653 and r_s_ = 0.765; n = 42), while there was significant correlation between qHF183 and *E. coli* and AllBac, but not Enterococci (r_s_ = 0.549, r_s_ = 0.734; n = 37) in DRG3 (Table S2). Although high levels of the human marker were detected in SWN, moderate correlation coefficients were observed between this marker and *E. coli* and AllBac (r_s_ = 0.423, r_s_ = 0.525; n = 42) suggesting that other factors determine the fecal pollution pattern at this site. Although low levels of qHF183 were observed in KMG1 there is a moderate relationship with *E. coli*, Enterococci and AllBac (r_s_ = 0.569; r_s_ = 0.547 and r_s_ = 0.430; n = 24).

The ruminant marker was moderately correlated with *E. coli*, Enterococci and AllBac in DRG2 (r_s_ = 0.600; r_s_ = 0.469, and r_s_ = 0.393, n = 40) and KGH (r_s_ = 0.552; r_s_ = 0.493 and r_s_ = 0.356; n = 42). In CBR, SWN and GCR a correlation with just AllBac was detected (r_s_ = 0.415, n = 42; r_s_ = 0.503, n=42; r_s_ = 0.552, n=18) (Table S2).

### Seasonality of MST markers and fecal contamination

To assess whether water quality was subject to seasonal variations, the dataset was grouped according to seasons: spring (March to May), summer (June to August), autumn (September to November) and winter (December to February) [21]. Using One Way Anova analysis, seasonal influence was detected for *E. coli* (P <0.001), Enterococci (P <0.001) and general *Bacteroidales* (P = 0.044), with the highest levels occurring during summer and autumn. The presence of the human marker (qHF183) was not influenced by the season during which the samples were taken (P = 0.719). In contrast, there was a strong seasonal effect on the levels of the ruminant marker (qCF128; P <0.001). The majority of samples testing positive for the ruminant marker in the upstream reaches of the Dargle (DRG1), and in the Kilmacanogue and Killough rivers, were taken in summer and autumn, with only a small percentage testing positive in winter (Table S3).

### The relationship between turbidity, FIB and MST markers

In general, there was a significant correlation between the turbidity of water samples and rainfall occurring on the day of sampling r_s_ = 0.410, the day before sampling r_s_ = 0.418 and both days r_s_ = 0.492), with a significant correlation at each sampling station (Table S4). With exception of CBR, all river sampling sites showed a significant correlation between rainfall and the levels of either *E. coli*, Enterococci, or both (Table S5). For the MST markers a positive correlation between the rainfall levels was observed with qCF128 in DRG2 and CBR. In contrast, a negative correlation of qHF183 with rainfall was observed in DRG3 and SWN.

A strong correlation between turbidity and FIB and MST was not observed. *E. coli* displayed a positive relation with turbidity in KMG1, qCF128 in DRG2 and CBR, whereas a negative correlation was observed with qHF183 in DRG1, DRG3 and SWN (Table S5).

## DISCUSSION

Several microbial source tracking markers have been tested successfully around the world [15,19,30,35,36]. However, selecting suitable microbial source tracking markers may be difficult since considerable variability may exists in the composition of the microbiome of individuals due to age, genetics, diet, health status or geographical location [14,37–39]. In this study we tested eight MST markers using end-point PCR and four markers using qPCR, with a view to determine their usefulness in their application for an effective management of water quality. The selected MST markers have been described elsewhere [4,9,24,25,28,40].

The variability of genetic marker abundance and prevalence in populations from other geographical locations suggests that the use of MST markers developed in a geographical area requires *a priori* characterization of the assay performance at each watershed of interest before being implemented [15,41–43]. Thus, in the first instance the selected MST markers were tested in fecal samples from known sources. All the PCR markers showed a high sensitivity and specificity, although none of them achieved 100% for both parameters. Although some of the MST markers were detected in hosts other than the intended ones, their abundance in the target group was always significantly higher than in the non-target hosts, demonstrating their suitability to distinguish between sources of pollution. These MST markers are diluted once the fecal matter is introduced in the environment. The lower levels reported in non-targeted samples are therefore likely to be below the detection limit once the fecal matter has entered the watershed.

The levels of the human marker (HF183) was variable in individual human stool samples. However, when the marker was tested in raw and treated sewage samples, 100% sensitivity and fairly constant marker levels were observed, which is in agreement with earlier observations by Seurinck et al [28]. These authors report levels of 10^5^ to 10^9^ gc gr^−1^ of wet human feces and 10^9^ to 10^10^ gc L^−1^ of sewage, similarly to the results of this study. The human gut harbors three predominant enterotypes dominated either by *Bacteroides*, *Prevotella* or *Ruminococcus* which could be related to for example diet [44–46], which might explain the high degree of variability in marker levels among individuals. However, the main sources of human pollution are sewer misconnection, effluents from WWTP or CSO, as well as septic tanks, rather than from individuals. The two human markers tested in this work: HF183 using end-point PCR and with SYBRGreen for qPCR were classified as the most sensitive and specific using binary analysis in a multi laboratory method evaluation study [47].

The effect of diet on the composition of the microbiota has also been reported for cattle, suggesting a strong correlation between bacterial microbiota structures and feeding practices. *Prevotella* spp. were the dominant group when animals were fed with unprocessed grain, while Ruminococcacea were the main group when other feeds were used [14]. In this study we tested the ruminant marker CF128 by end-point PCR [4] and since it showed a good specificity, a SYBRGreen assay with this primer was developed. The probe-labeled assay targeting the cow specific marker CowM3 [29] was used to be able to discern among cattle and other ruminants (especially sheep and deer). The CF183 marker was detected in cows, sheep, deer and goat by PCR and qPCR and showed a cross reaction mainly with horse and pig. However, it was four orders of magnitude more abundant in ruminant than non-ruminant samples. The concentration of CowM3 marker was around two orders of magnitude lower than CF183. This marker targets the gene encoding sialic acid-specific 9-*O*-acetylesterase secretory protein. Whereas a single copy of this gene is present in the genome, up to 7 copies of the 16S rRNA gene per genome have been reported in several *Bacteroidales* species [48]; the latter is therefore easier to detect following dilution of fecal matter in the water column. In contrast to previous reports, 50% of the sheep samples in the current study tested positive for the CowM3 marker, with similar levels of the marker in cow and sheep feces. In contrast, all deer fecal samples tested negative for this marker. Dorai-Raj et al [49] showed that bovine and ovine microbiotas are very similar, preventing the development of a bacterial MST marker to differentiate between these species. Sheep and cows are kept in intensive farming in many countries, whereas in Ireland these animals are mainly kept on pastures, which may account for the similarities in microbiota and poor specificity of the CowM3 marker. This problem may be solved using markers targeting mitochondrial DNA [25,50,51].

In this study, markers targeting mitochondrial DNA from pig and sheep showed a high specificity and sensitivity (Table 2). Just a few positives were observed in human fecal samples, raw and treated sewage, which might be due to a recent ingestion of pork and sheep products. Other studies analyzing mitochondrial DNA detected the bovine marker in individuals who had eaten beef 24 hours before the sample collection; however the levels were two orders of magnitude lower than those of the human marker [51]. In this study we did not perform a quantitative analysis for these markers which might have shown much lower concentration of pig or sheep mitochondrial DNA in human feces. Therefore, mitochondrial DNA may prove to be a good option to be used with a combination of bacterial markers to help to discriminate among animals with similar microbiota such as cow, sheep and deer [50,52].

Two *Bacteroidales* pig markers were also evaluated in this work, whereas PF163 showed a higher sensitivity than Pig-2-Bac, a higher specificity was achieved by the latter, suggesting the use of a combination of markers to increase source tracking resolution [53]. Finally the gull marker showed a very high specificity, not being detected in other avian samples like swans or peacocks.

The HF183 and CF183 markers were evaluated in several European countries, including Ireland [30], which showed 88% sensitivity and 100% specificity for HF183 and 100% sensitivity and 96% specificity for CF183, which is slightly higher than what was observed in this study. Both markers were also evaluated in Galway (West of Ireland) what allows us to compare regional geographic variability [54]. For HF183 a sensitivity of 12% for individual fecal samples and 70% for sewage was reported with 100% of specificity, whereas 94% sensitivity and 95% specificity was observed for CF183. In the latter cross-reaction was observed with mainly pig fecal samples, as is the case in this study. These markers therefore perform in a consistent manner in different areas, and at different times.

The levels of the host specific *Bacteroidales* markers (qCF128) in the target host were several orders of magnitude higher, ranging from 10^7^-10^8^ gc 100 mg wet wt^−1^, than in other hosts, similar to what was observed previously using samples taken in California [55]. The very high levels of these markers in their target hosts mean that remain above the detection limit following release in a waterbody.

The next step was to evaluate the performance of the selected MST markers in a relatively small catchment with well-defined land uses. The analysis of land uses in the watershed, the visual inspection of the area, together with the microbial source tracking analysis allowed a very accurate assessment of the main hazards and sources of pollution in the catchment. The Dargle catchment has a population of 58,745 (census of 2006). This study showed that human fecal pollution is the main impact in the catchment. It has been observed that human pollution is mainly coming from the effluent of a WWTP (between DRG2 and DRG3), a non-detected sewer misconnection/s in the Kilmacongue River (KMG1 and KMG2) and the Swan River crossing the biggest urban area in the catchment showing even higher levels than those observed in treated sewage. Human fecal pollution was also prevalent with lower levels in agricultural areas probably due to leaks of septic tanks from the farms and houses spread around. A study combined land use information and small stream sampling reported a positive significant correlation between abundance of human associated markers and septic systems following a wet weather event [19]. Unfortunately in this study we did not have information about septic systems and sanitary sewer lines locations which may strongly support the success of the MST strategies. The levels of the human associated maker in this study showed a negative significant correlation in DRG3 and SWN with rainfall suggesting a dilution of the pollution after rainfall events. There were no differences among seasons between the levels of human marker, indicating human pollution as a constant pressure along the year.

The presence of the CF128 marker was higher in areas surrounded by agriculture and the levels were fairly even along the catchment being less prevalent in urban areas (Swan River) showing a basal ruminant pollution in the river. Data from the number of animals was obtained from the Census of Agriculture 2000 from the Central Statistics Office Ireland and reported around 3,280 cattle and 24,115 sheep heads in the catchment. Unfortunately, this data is reported for electoral districts, instead of the catchments areas used in this study, therefore animal density could not be used to match the land uses with GIS. However these numbers together with wild deer in the area might explain this basal fecal contamination. Significant correlation between the levels of the ruminant associated marker with *E. coli*, Enterococci in DRG2 and KGH indicates ruminant pollution as the main pressure in these points. The high levels of ruminant associated marker in CBR and its relation with AllBac, but not with FIB, suggests an additional source of pollution in this point such as bird pollution holding reduced numbers of *Bacteroidales* such it has been observed in gulls and swans. The ruminant marker showed seasonal patterns being more abundant in summer and autumn and a positive correlation with maximum temperature in DRG1, DRG3, KGH and GCU, four sampling sites surrounded by forest and agriculture. Since lower persistence of *Bacteroidales* markers have been reported with higher temperatures [56], the higher levels of the ruminant marker may be related to a higher contribution in warm seasons. Additionally, in two points (DRG2 and CBR) surrounded by agriculture the ruminant marker correlated with rainfall and turbidity, suggesting an important input coming from run-off. Identifying these inputs to the system can improve catchment management establishing agricultural best management practices (BMP) like cattle exclusion fencing or controlled tile drained management [57,58].

At most sites, the *Bacteroidales* marker showed a high correlation with *E. coli* and Enterococci, while a low correlation was observed high in upstream waters. Here, none or very low concentrations of *E. coli* but high levels of the *Bacteroidales* marker was detected. . This suggests that the marker may be detecting non-intestinal environmental bacteria, in accordance with other studies detecting *Bacteroidales* markers in pristine and drinking waters [59,60]. Its use as fecal indicator may depend of the sampling site and the levels of pollution [61]. *E. coli* and Enterococci in general showed a high seasonal pattern and high correlation with maximum temperature detecting higher levels in warmer periods. A lower persistence of these markers with higher temperature has also extensively described [62–64] suggesting that the high levels are mainly related to major pollution inputs during such periods. Moreover, there was a positive correlation with FIB and rainfall, with the exception of DRG3 where a negative correlation was found due to the dilution of the source of the pollution (the WWTP effluent) and CBR where no correlation was found. In the case of the MST markers analyzed in this study, such a correlation was only observed at a few sites, suggesting a potential of other fecal input in the catchment with rainfall events. These results confirm that BMP of run-off could improve water quality.

The MST toolbox evaluated in this study shows a good potential to be used for a better management and assessment of fecal pollution. However, it is advisable to evaluate the markers on fecal samples collected in the area where they will be applied in order to know their specificity, sensitivity, prevalence and understand their patterns in the environmental samples. The combined use of FIB, MST markers, environmental data and Geographical Information System to integrate land uses analysis with visual exploration, achieves appreciably enhanced description of the potential main causes of fecal pollution in a river catchment area. This data may be applied to develop hydrological models integrating bacterial data to facilitate the application of measures to eliminate or reduce the levels of fecal pollution at the source.

## ACKNOWLEDGMENTS

We thank Jonathan Sexton (Wicklow County Council) for his advice and assistance in collecting samples, Our Lady’s Children’s Hospital,Crumlin, UCD Veterinary Hospital, Lyons UCD farm and Mr. Daragh La Grue who assisted in fecal sample collection. This project was funded by a grant from the Irish Environmental Protection Agency under the Science, Technology, Research & Innovation for the Environment (STRIVE) Programme 2007-2013.

